# Pain-related opioidergic and dopaminergic neurotransmission: Dual Meta-Analyses of PET Radioligand Studies

**DOI:** 10.1101/2020.09.18.302943

**Authors:** Sergio Guerra Garcia, Andrea Spadoni, Jennifer Mitchell, Irina A. Strigo

## Abstract

Molecular mechanisms of the interaction between pain and reward associated with pain relief processes in the human brain are still incompletely understood. This is partially due to the invasive nature of the available techniques to visualize and measure metabolic activity. Positron Emission Tomography (PET) radioligand studies using radioactive substances are still the only available modality to date that allows for the investigation of the molecular mechanisms in the human brain. For pain and reward studies, the most commonly studied PET radiotracers are [^11^C]-carfentanil (CFN) and [^11^C]- or [^18^F]-diprenorphine (DPN), which bind to opioid receptors, and [^11^C]-raclopride (RAC) and [^18^F]-fallypride (FAL) tracers, which bind to dopamine receptors. The current meta-analysis looks at 15 pain-related studies using opioid radioligands and 8 studies using dopamine radioligands in an effort to consolidate the available data into the most likely activated regions. Our primary goal was to identify regions of shared opioid/dopamine neurotransmission during pain-related experiences. SDM analysis of previously published voxel coordinate data showed that opioidergic activations were strongest in the bilateral caudate, thalamus, right putamen, cingulate gyrus, midbrain, inferior frontal gyrus, and left superior temporal gyrus. The dopaminergic studies showed that the bilateral caudate, thalamus, right putamen, cingulate gyrus, and left putamen had the highest activations. We were able to see a clear overlap between opioid and dopamine activations in a majority of the regions during pain-related processing, though there were some unique areas of dopaminergic activation such as the left putamen. Regions unique to opioidergic activation include the midbrain, inferior frontal gyrus, and left superior temporal gyrus. By investigating the regions of dopaminergic and opioidergic activation, we can potentially provide more targeted treatment to these sets of receptors in patients with pain conditions. These findings could eventually assist in the development of more targeted medication in order to help treat pain conditions and simultaneously prevent physical dependency.

## 1. Introduction

Pain conditions are widespread, affecting around 20% of the US population, and are often treated with opioid medications (Dahlhamer et al., 2018). However, recently there has been a push to reduce opioid use, as such medications often lead to addiction (Cowan et al., 2003; Fishbain et al., 1992). Incidents of fatal opioid overdoses have now reached epidemic proportions, accounting for two thirds of all drug-related deaths in the United States (Wilson, 2020). Prescription opioids in particular remain a primary driver of opioid-related deaths (McLachlan, Hay et al. 1994). Notably, consumption of exogenous opioids lead to changes within endogenous opioid and dopamine neurotransmission (Kosten and George 2002), both of which play fundamental roles in pain processing, pain modulation, and pain relief (Fields 2018). Investigating whether dopaminergic and opioidergic activity are released in the same regions during similar conditions may enlighten us on how it is they work together to create and process the perception of pain/relief.

Recent advances in non-invasive neuroimaging have allowed for the exciting investigation of human brain function and structure with respect to human pain processing (Tracey and Mantyh 2007). Functional MRI studies have allowed us to observe how various brain regions are influenced by experimental and clinical pain conditions. Among these, the insular cortex is activated in a wide variety of pain paradigms as well as in emotional and interoceptive aspects of pain (Jensen et al., 2016). The brain stem and anterior cingulate cortex provide connectivity among brain areas related to the subjective perception and modulation of pain (Ploner et al. 2010). The ventral striatal area, specifically the nucleus accumbens (NAc) is part of the reward system that modulates dopamine signaling in response to a pain stimulus, the activation of this area occurs with both acute pain and pain relief (Becerra et al. 2013). The PFC has been widely implicated in the neuromodulation of pain through its connections with areas such as the cerebral neocortex, hippocampus, periaqueductal grey (PAG), thalamus, amygdala, and basal nuclei (Ong, Stohler, and Herr 2019).

Nevertheless, the only method by which to measure neurotransmission in the human brain is still Positron Emission Tomography (PET). The endogenous opioidergic system is normally assessed using [^11^C]-carfentanil (CFN) and [^11^C]- or [^18^F]-diprenorphine (DPN) as radiotracers. CFN is a selective agonist at the μ-opioid receptor (MOR), which is thought to preferentially bind to the μ1 receptor sub-type (Eriksson and Antoni 2015), whereas DPN is a weak partial agonist with equal affinity for the MOR, κ-opioid receptor (KOR), and δ-opioid receptor (DOR). The endogenous dopaminergic system is normally assessed by [^11^C]-raclopride (RAC) and [^18^F]- fallypride (FAL) tracers, which are selective antagonists on D2/D3 dopamine receptors.

Information about both basal levels of receptor availability and changes in availability caused by alterations in endogenous dopamine and opioid concentrations can be evaluated, as these radioligands are sensitive to endogenous opioid and dopamine levels. PET radioligand studies have provided valuable information on the idiosyncratic concentrations and distributions of such neurochemicals across the brain, and researchers have used these data to better understand sensory and emotional pain processes (e.g. (Zubieta, Smith et al. 2001), as well as addiction processes associated with opioids (Greenwald, Johanson et al. 2003) and dopamine (Volkow, Fowler et al. 2009). Notably, as demonstrated by prior animal studies (Benarroch 2012), there is anatomical overlap of neurotransmitter systems. Studies demonstrating such overlap of these neurochemicals in the human brain are limited (Goldman-Rakic, Lidow et al. 1990) and one of the major limitations of current PET technology is that it only allows for the imaging of a single radiotracer at a time. Recent research has provided insight into the fact that endogenous opioids are released in the ventral striatum, insula, anterior cingulate cortex (ACC), prefrontal cortex (PFC), and brain stem during pain experience, though knowing whether dopamine is also released in many of the same regions during similar pain-related conditions may demonstrate that the two neurotransmitter systems might be working together to create and process the perception of pain/relief. Thus, an increased understanding of the brain regions where these neurotransmitters co-localize in the human brain may further elucidate pain processing and pain behaviors. The present dual meta-analysis specifically aims to create a map of the opioid and dopamine neurotransmitter systems in the brain throughout painful (both sensory and emotional) conditions. An understanding of the brain regions where these neurochemicals either uniquely concentrate or overlap may assist in the development of less potent and more targeted medications in order to help treat pain conditions while simultaneously preventing physical dependence.

### 1.1 Neural Correlates of Opioid Receptors

The endogenous opioid system consists of 3 families of opioid peptides: β-endorphin, enkephalins, and dynorphins, and 3 families of receptors: μ (MOR), δ (DOR), and κ (KOR) (reviewed in detail elsewhere e.g. (Benarroch 2012). Though endogenous opioids can be found extensively throughout the central and peripheral nervous system, they are particularly concentrated in circuits involved in pain modulation, reward, responses to stress, and autonomic control. Some of the areas with the highest opioidergic receptor concentration are the cerebral cortex, brainstem, thalamus, striatum, hypothalamus, hippocampus, and dorsal horn (Benarroch 2012). Radioligand studies of opioidergic receptors included in this meta-analysis have reported opioid activations in a number of cortical and subcortical regions, including frontal cortices (Jones, Kitchen et al. 1999, Willoch, Schindler et al. 2004, Dougherty, Kong et al. 2008, Klega, Eberle et al. 2010, Mueller, Klega et al. 2010, Maarrawi, Peyron et al. 2013, Wey, Catana et al. 2014), insula (Jones, Kitchen et al. 1999, Willoch, Schindler et al. 2004, Dougherty, Kong et al. 2008, Mueller, Klega et al. 2010, Wey, Catana et al. 2014, Brown, Matthews et al. 2015), anterior cingulate cortex (Jones, Kitchen et al. 1999, Willoch, Schindler et al. 2004, Sprenger, Valet et al. 2006, Dougherty, Kong et al. 2008, Mueller, Klega et al. 2010, Maarrawi, Peyron et al. 2013, Wey, Catana et al. 2014), thalamus (Jones, Kitchen et al. 1999, Willoch, Schindler et al. 2004, Dougherty, Kong et al. 2008, Wey, Catana et al. 2014, Brown, Matthews et al. 2015) and putamen (Jones, Kitchen et al. 1999, Wey, Catana et al. 2014, Brown, Matthews et al. 2015).

### 1.2 Neural Correlates of Dopamine Receptors

Dopamine is widely involved in circuits encoding reward, aversion, salience, uncertainty, novelty (Bromberg-Martin, Matsumoto et al. 2010), and pain modulation (Wood 2008). The midbrain houses some of the areas with the highest dopaminergic concentrations, namely the Ventral Tegmental Area (VTA) and Subtantia Nigra (SN), as well as projections that lead to the dorsal striatum, nucleus accumbens, amygdala, hippocampus, and prefrontal cortex (PFC).

Radioligand studies of dopaminergic receptors have reported activations in a number of cortical and subcortical regions, including bilateral putamen (Wood, Schweinhardt et al. 2007, Scott, Stohler et al. 2008, Berman, Hallett et al. 2013), left putamen (Martikainen, Nuechterlein et al. 2015), left caudate (Scott, Stohler et al. 2007, Wood, Schweinhardt et al. 2007, Scott, Stohler et al. 2008, Berman, Hallett et al. 2013, Martikainen, Nuechterlein et al. 2015), right caudate nucleus (Wood, Schweinhardt et al. 2007, Scott, Stohler et al. 2008, Martikainen, Nuechterlein et al. 2015), right nucleus accumbens (Scott, Stohler et al. 2008, Martikainen, Nuechterlein et al. 2015, Pecina, Love et al. 2015), left nucleus accumbens (Scott, Stohler et al. 2008), and globus pallidus (Wood, Schweinhardt et al. 2007).

### 1.3 Interaction between opioidergic and dopaminergic neurotransmission

In many regions, opioids are co-expressed with other neurotransmitters (Benarroch 2012). Specifically, opioid neurotransmitter systems interact with dopamine not only in the midbrain but also in projection areas such as the striatum, a key area implicated in reward and pain relief (Fields 2018). In addition to its established role in the reward system, many studies have suggested that the NAc also serves as a meeting point for multiple components of pain processing and analgesia. Some of the regions connected by the NAc include the ACC, PFC, thalamus, amygdala, somatosensory cortex, and the spinal cord, which span the reward and pain systems (Harris, 2020). Striatal medium spiny neurons express both dopamine and opioid receptors (Pollard, Llorens-Cortes et al. 1977, Ambrose, Unterwald et al. 2004). Blocking striatal opioid receptors leads to attenuated amphetamine-induced locomotion and impulsivity (Gonzalez-Nicolini, Berglind et al. 2003, Wiskerke, Schetters et al. 2011), whereas dopamine D2 receptor (DRD2) blockade inhibits the rewarding effects of morphine in opiate dependent rats (Laviolette, Nader et al. 2002). Likewise, activating μ-opioid receptors (MOR) modulates the mesolimbic dopamine system. As shown in rats, morphine modulates the release of dopamine by disinhibition through GABAergic interneurons in the midbrain ventral tegmental area (VTA) (Jalabert, Bourdy et al. 2011). In humans, alfentanil triggers dopamine release in the striatum (Hagelberg, Kajander et al. 2002). Likewise, activating D2/D3 receptors modulate mesolimbic opioid system. As shown in several studies in humans, amphetamine releases endogenous opioids in the striatum, as well as in insular and anterior cingulate cortices (Colasanti, Searle et al. 2012, Mick, Myers et al. 2014), confirming the interdependence of the two systems. To date only a few studies have examined the overlap between opioidergic and dopaminergic receptor binding in the same individual, though there is data showing overlap in the ventral striatum and dorsal caudate nucleus (Tuominen, Tuulari et al. 2015; Scott et al., 2007; Karjalainen et al., 2017; Winterdahl et al., 2019; DaSilva et al., 2019).

From current literature it is evident that there is high variability across PET radioligand studies regarding the foci of activations. These inconsistencies may be attributed to several factors including, but not limited to, demographic characteristics of the sample such as age and sex, the presence of comorbid disorders and/or childhood adversity, differences in protocols and processing, and statistical analyses. Here we applied seed-based d mapping meta-analyses (Radua, Mataix-Cols et al. 2012) that have been established as a standard tool for identifying coordinate based convergence of voxel based imaging data. This analysis was conducted separately for the opioidergic radioligands and for the dopaminergic radioligands and enabled us to explore overlap between the two neurotransmitter systems.

## 2. Methods

### 2.1 Article Selection

We conducted a literature search for radioligand PET studies that were published between June 1999 and September 2017 using several sources including Pubmed (https://www.ncbi.nlm.nih.gov/pubmed/) and Embase (https://www.embase.com) to find reports with the radioligands of interest such as “carfentanil”, “diprenorphine”, “raclopride”, “fallypride”. The study Flow Chart is depicted in **Figure 1**. The final article search was conducted on July 2019 and yielded 393 PET imaging studies using the radioligands of interest to be considered for further review. The studies included in our analysis mapped brain regions associated with opioidergic and dopaminergic receptors using CFN, DPN, RAC and FAL radioligands. We investigated general opioid receptors (MOR, DOR, and KOR) via diprenorphine (DPN) while selective radioligands like carfentanil (CAR) was used to study MOR. Both fallypride (FAL) and raclopride (RAC) were used to study dopamine type-2 and type-3 (D2/D3) receptors.

**Fig.1.**
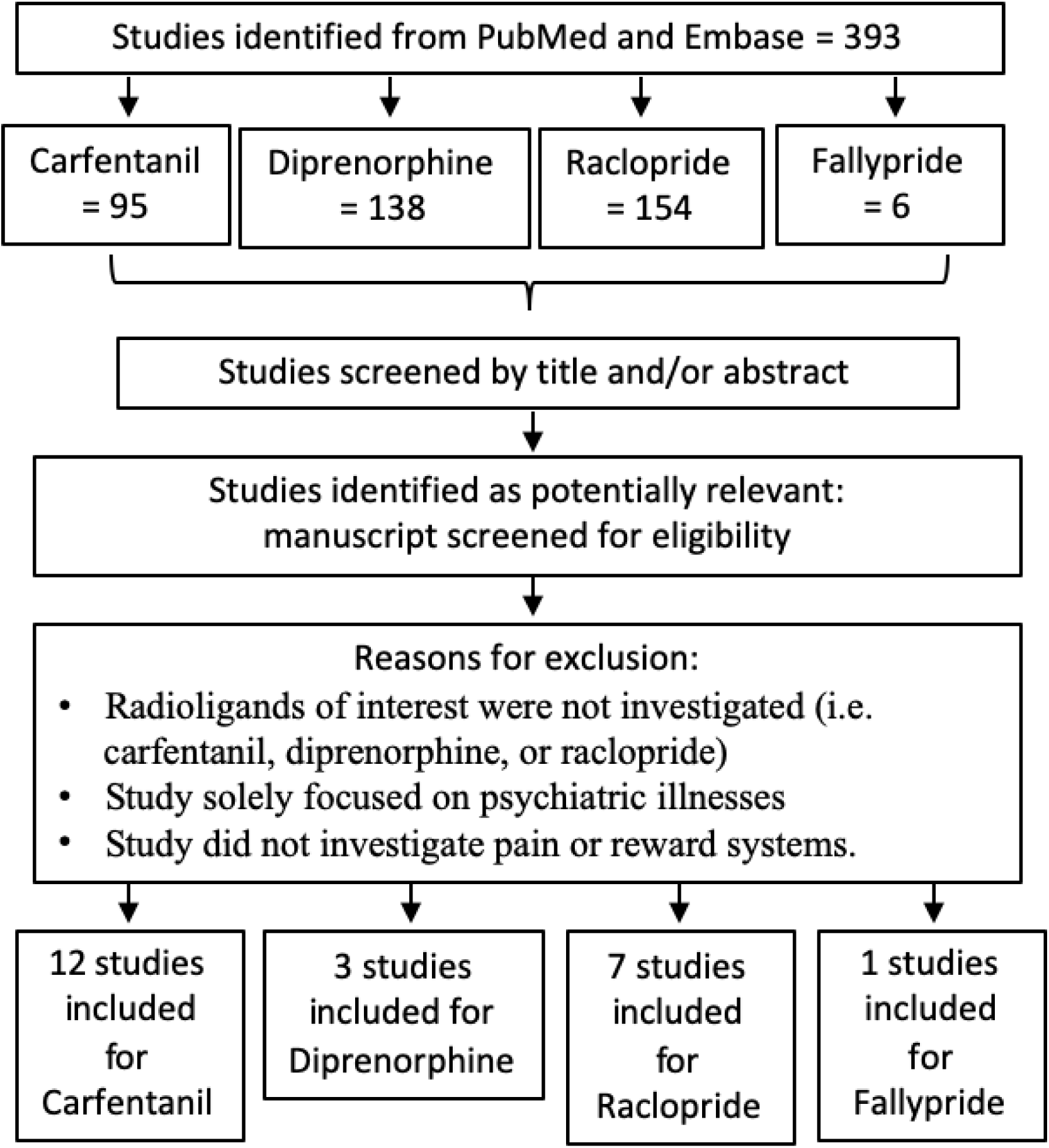
Flow chart of search strategy and study selection for meta-analysis. Study search and screening procedures repeated for PubMed and Science Direct

All studies of pain-related processing and experiences that provided voxel-based coordinates of the observed peak activations were included. Due to the variable nature of PET radioligand studies, i.e., examining acute and chronic pain conditions as well as correlational changes, we limited our inclusion to studies only reporting within-group observations, i.e., only those studies that reported changes in endogenous dopaminergic or opioidergic transmission as a result of experimental pain or endogenous chronic pain manipulation (painful state vs. non-pain state). Studies reporting between-group observations (i.e., changes in the endogenous neurotransmission between healthy subjects and individuals with chronic pain were excluded from the meta-analyses.

### 2.2 Inclusion/exclusion criteria for activation foci

Significant peak activation coordinates and corresponding intensity values for each of the articles in **Table 1** were extracted for differences in experimental conditions. If studies included two separate control conditions, only foci from one of the within-subject comparisons were used to avoid using foci from the same participant twice. In these cases, the selected contrast compared a painful or unpleasant condition with a non-painful condition. If a study conducted a whole-brain and a region of interest (ROI)-based analysis [Brown, Martikainen, Peciña, Harris, Scott 2007, Berman], coordinates from both analyses were included separately, provided that the ROIs were not reported in the whole-brain results.

**Table 1:** Characteristics of studies included in the meta-analysis

| Study | Sample | Stimulation | Sex (M,F) | N | Age | Type | Significance |
| --- | --- | --- | --- | --- | --- | --- | --- |
| **Opioid** |  |  |  |  |  |  |  |
| **Carfentanil** |  |  |  |  |  |  |  |
| Prossin, 2015 | Healthy controls | Masseter muscle pain | 12, 22 | 34 | 22±11 | WBA | p<0.001 |
| Zubieta, 2001 | Healthy controls | Masseter muscle pain | 13, 7 | 20 | 24±2 | WB | p<0.05 |
| Scott, 2007 | Healthy controls | Masseter muscle pain | 14, 0 | 15 | 27±5 | WBA & ROI | p < 0.01 |
| Wager, 2007 | Healthy controls | Thermal Pain | n/a | 15 | 25±5 | ROI | p < 0.001 |
| Scott, 2008 | Healthy controls | Masseter muscle pain | 9, 11 | 20 | 24±3 | WBA | p < 0.0001 |
| DaSilva, 2014 | Migraine | Headache pain | 3, 4 | 12 | n/a | WBA | n/a |
| Nascimento, 2014 | Migraine | Thermal Pain | 3, 3 | 6 | 26 | WBA | p < 0.0001 |
| Pecina, 2015 | Healthy controls | Masseter muscle pain | 21, 29 | 50 | 26±0.7 | WBA | p < 0.0001 |
| Zubieta, 2003 | Healthy controls | Sad mood induction | 0, 14 | 14 | 36±9 | WBA | P < 0.001 |
| Hsu, 2013 | Healthy controls | Rejection Pain | 5, 13 | 18 | 32±12 | VOI | p < 0.05; p < 0.01 |
| Harris, 2009 | Fibromyalgia | Fibromyalgia pain | 0, 20 | 20 | 44.3±13.6 | WBA & ROI | P < 0.001 |
| Pecina, 2013 | Healthy controls | Masseter muscle pain | 19, 18 | 37 | 30±10 | WBA & ROI | p < 0.001; p < 0.05 |
| **Diprenorphine** |  |  |  |  |  |  |  |
| Maarrawi, 2007 | Neuropathic pain | Chronic pain | 4, 4 | 8 | 55.2±11 | ROI | p < 0.0005 |
| Wey, 2014 | Healthy controls | Pressure pain | 4, 4 | 16 | 24 | ROI | p < 0.05 |
| Jones, 1999 | Trigeminal Neuralgia | Neuralgia pain | 6, 0 | 6 | 61.5 ± 9.5 | ROI | p < 0.01 |
| **Dopamine** |  |  |  |  |  |  |  |
| **Raclopride** |  |  |  |  |  |  |  |
| Berman, 2013 | Healthy controls | Writer's cramp pain | 10, 5 | 15 | 52.4 ± 9 | WBA & ROI | p < 0.05 |
| Martikainen, 2015 | Healthy controls | Masseter muscle pain | 18, 14 | 16 | 35 ± 11 | WBA & ROI | p < 0.05; p < 0.005 |
| Pecina, 2015 | Healthy controls | Masseter muscle pain | 21, 29 | 50 | 26±0.7 | WBA | p<0.05; p < 0.0001 |
| Scott, 2006 | Healthy controls | Masseter muscle pain | 18, 7 | 25 | 27±5 | WBA | p<0.0001; p<0.01 |
| Scott, 2007 | Healthy controls | Masseter muscle pain | 14, 0 | 14 | 27±5 | ROI | P < 0.01 |
| Scott, 2008 | Healthy controls | Masseter muscle pain | 9, 11 | 20 | 24±3 | WBA & ROI | p = 0.05, p<0.0001 |
| Wood, 2007 | Healthy controls | Tibialis muscle pain | 0, 22 | 11 | n/a | ROI | p < 0.05 |
| **Fallypride** |  |  |  |  |  |  |  |
| Jarcho, 2015 | Healthy controls | Thermal Pain | 0, 15 | 15 | 24±3.11 | WBA | p = 0.001 |

WBA = Whole brain analysis; VOI = Volume of interest; ROI = Region of interest

Twenty-three radioligand studies (**Table 1**) were included in the current investigation based on the inclusion and exclusion criteria, and produced a total of 451 subjects from which peak voxels and clusters were extracted and transformed into Montreal Neurological Institute (MNI) space if they were provided in Talairach space. Of the 393 articles found, 368 were excluded for the following reasons: 1) Radioligands of interest were not investigated (i.e. CFN, DPN, RAC or FAL), 2) study solely focused on how neurological disorders affect the neurotransmitter of interest (e.g. Parkinson’s Disease), 3) study did not investigate pain or reward systems; 4) study provided only correlational relationship with the baseline neurotransmitter release; and 5) studies only reported on the between-group comparisons between healthy controls and individuals with chronic pain without within-subject experimental manipulation.

### 2.3 Meta-analysis

We conducted a coordinate-based random-effects meta-analysis using sdmGUI software version 5.142 (https://www.sdmproject.com/). Coordinates reported in Talairach space were converted to MNI space using the “convert peaks” tool provided by sdmGUI. We also gathered the peak intensity for each coordinate in the form of t-scores, otherwise they were converted from z-scores or p-values into t-scores using the “Convert peaks” feature of sdmGUI. Twelve studies were identified that examined mu-opioid receptors using CFN and 3 studies were identified that assayed all opioid receptors using DPN; these 15 studies were included in the opioidergic meta-analysis. Seven studies were used to examine D2/D3 receptors using RAC and 1 study used FAL radioligand; these 8 studies were included in the dopaminergic meta-analysis.

An anisotropic effect-size-based seed-based d mapping (AES-SDM) statistical analysis was conducted to determine the mean regions of the brain that were consistently activated. Coordinates from each of the radioligands were first analyzed separately to examine the neural regions involved across each of the neurotransmitters. In order to determine the robustness of each of the studies, a jackknife analysis available through sdmGUI was conducted. Analyses from opioidergic and dopaminergic radioligands were then combined to examine the spatial overlap and non-overlap between mu-opioid, general opioid, and D2/D3 receptors using AFNI function *3dcalc* (Cox 1996).

When there were differences in the thresholding in whole-brain and ROI analyses, they were entered separately in order to correct for the threshold differences conducted by the authors. The areas of activation found in the results were shown in high consistency throughout the included studies and the z-scores indicate the likeliness of activation in these areas. Furthermore, a jack-knife analysis was also conducted to account for voxels with only a few studies contributing, this way single studies that could possibly dominate the results would not skew the meta-analysis (Cutler et al., 2018). False Discovery Rate (FDR) threshold was set to p < 0.005 in order to assess for random spatial associations between experiments. The standard AES-SDM thresholds (uncorrected voxelwise p-value of p < 0.005, extent threshold clusters for ≥ 10 voxels, and z values of greater than or equal to Z ≥ 1, which are proposed to optimally balance sensitivity and specificity) were used.

## 3. Results

### 3.1 Opioid

To increase power, we combined studies that evaluated selective (μOR) and non-selective (μOR, δOR, and κOR) opioid receptor radioligands. This resulted in a total of 15 articles, with a total n = 283 subjects, yielding 121 foci. When all opioidergic activity was combined, 3 clusters (**Table 2**) were observed with peak activations in the right striatum, cingulate gyrus extending to supplementary motor area, and midbrain region of the brainstem. The largest cluster, with a maximum peak in the right striatum, is composed of 7840 voxel activations spreading across bilateral amygdalae, bilateral basal ganglia, bilateral insulae, subgenual cingulate and frontal pole regions (**Figure 2**).

**Table 2.** MNI coordinates for brain regions in mu-opioid activations through carfentanil

| *MNI coordinate* |  |  |  |  |  |  |
| --- | --- | --- | --- | --- | --- | --- |
| x | y | z | *SDM-Z* | *P value* | Cluster Size | Brain Region, Brodmann Area |
| 20 | 4 | -6 | 4.76 | ~0 | 6588 | Right striatum |
| 2 | 8 | 46 | 2.99 | 0.00001 | 453 | Right supplementary motor area, BA 32 |
| -56 | -18 | 6 | 2.55 | 0.0002 | 268 | Left superior temporal gyrus, BA 48 |

*Threshold Voxel threshold: $p < 0.005$ . Peak height threshold: peak SDM-Z > 2. Extent threshold: cluster size $\geq 10$ voxels.*

**Figure 2:**
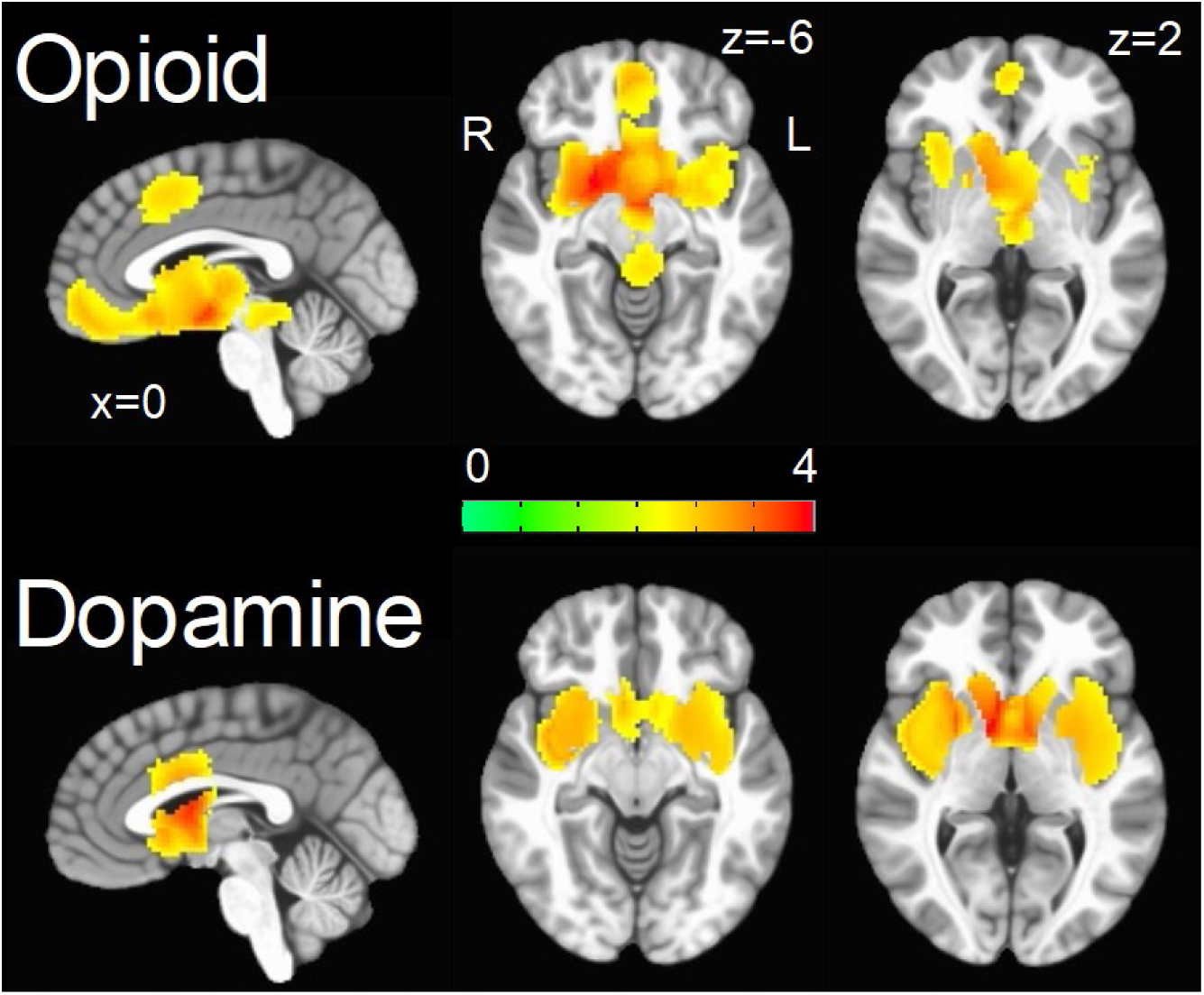
Areas of significant change in the endogenous opioid (top) and dopamine (bottom) transmission during pain-related experiences. Gradient bar represents SDM value.

Jackknife sensitivity analysis revealed that activations in the right caudate, right insula, and the midbrain, were highly robust, as they were replicated in all 15 studies. Areas in the right putamen were highly influenced by Pecina et al., 2014. Opioidergic activations in the inferior frontal gyrus were determined to not be robust, as there was a lack of activation in 7 of the 15 studies (**Table 4**).

**Table 3.** Peak MNI coordinates for brain regions in D2/D3 activations

| *MNI coordinate* |  |  |  |  |  |  |
| --- | --- | --- | --- | --- | --- | --- |
| x | y | z | SDM-Z | P value | Cluster Size | Brain Region |
| 1 | 3 | 2 | 3.816 | ~0 | 5320 | Righ Striatum |
| 0 | 9 | 29 | 2.574 | 0.0001 | 690 | Cingulate Gyrus (BA 24) |

*Threshold Voxel threshold: $p < 0.005$ . Peak height threshold: peak SDM-Z > 2.0. Extent threshold: cluster size $\geq 10$ voxels.*

**Table 4.**
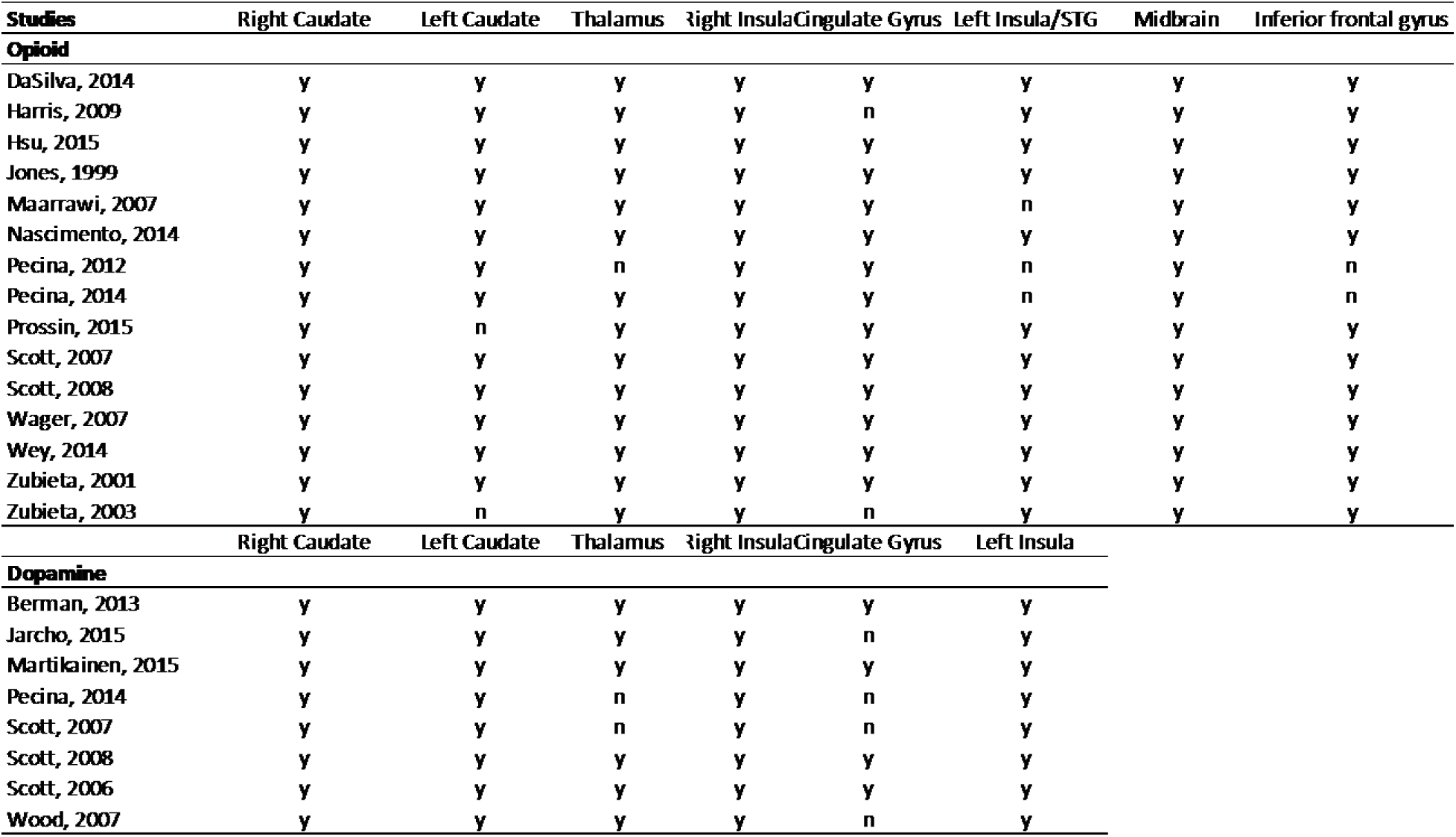
Jack-Knife sensitivity analysis for each significant cluster

### 3.2 Dopamine

Eight manuscripts evaluating dopaminergic activity via RAC and FAL reported a total of 49 foci with a total n = 166. Two clusters were observed with peak activations in the right striatum and cingulate gyrus, respectively (**Table 3**). The right striatum cluster included 5320 voxels and spread across bilateral insulae, thalamus, and basal ganglia regions (**Figure 2**, bottom).

Jackknife sensitivity analysis revealed that activations in the bilateral caudate and bilateral insulae were highly robust, as they were replicated in all eight studies. It should also be noted that activity in the thalamus was highly influenced by Pecina et al., 2014 and Scott et al., 2007. D2/D3 activations in the cingulate gyrus were determined to not be robust as there was a lack of activation in 4 of the studies (**Table 4**).

### 3.3 Overlap

In order to explore co-localization of endogenous opioid and dopaminergic neurotransmission during pain-related experiences, we created a conjunction between opioid and dopamine thresholded meta-analytical maps. The resulting map is displayed in **Figure 3**. We found biggest overlap within striatal regions followed by overlap within anterior cingulate (Table 5). The striatal cluster was large and spread to include bilateral amygdalae, bilateral anterior insulae, ventral cingulate, and bilateral caudate (**Figure 3**).

**Table 5.** Peak MNI coordinates for overlap brain regions of opioid and D2/D3 activations

|  |  |  |  |  |
| --- | --- | --- | --- | --- |
| MNI coordinate |  |  | Cluster Size | Brain Region |
| x | y | z |  |  |
| 3 | 6 | -2 | 3877 | Right Striatum |
| 1 | 15 | 34 | 58 | Right Cingulate Gyrus (BA 32) |

Extent threshold: cluster size ≥ 10 voxels.

**Figure 3:**
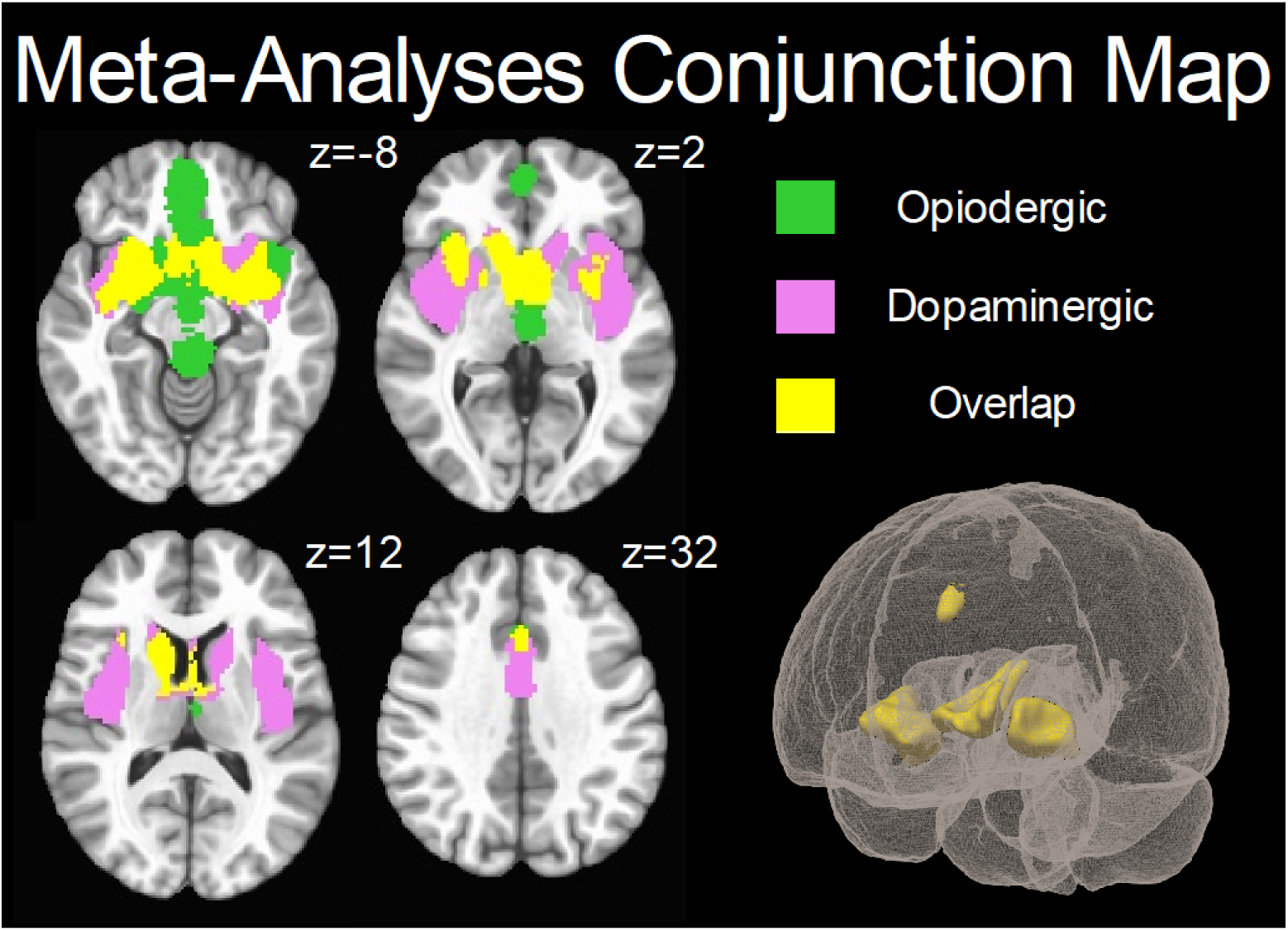
Areas of overlap (yellow) between endogeneous opioid (green) and dopamine (pink) transmission during pain-related experiences.

## 4. Discussion

The goal of this investigation was to provide the meta-analytical map of opioidergic and dopaminergic transmissions and their overlap through pain-related conditions including experimental pain, endogenous clinical pain, and emotional pain in humans using the opioidergic and dopaminergic radioligands CFN, DPN, RAC and FAL. Using the existing literature, we completed two comprehensive meta-analyses of PET radioligand activations using opioidergic and dopaminergic radioligands in sensory/emotional pain studies.

Consistent with the animal literature, we found sensory/emotional pain related opioidergic activation within striatum, cingulate gyrus extending to supplementary motor area and the midbrain region of the brainstem. The striatal cluster was large and spread across bilateral amygdalae, thalamus, bilateral insulae, subgenual cingulate and frontal pole regions. Our second meta-analysis of dopaminergic activation showed sensory/emotional pain-related dopaminergic activation within striatum and cingulate gyrus. The striatal cluster was large and spread across bilateral insulae and the thalamus.

Our primary goal was to identify regions of shared opioid/dopamine neurotransmission during pain-related experiences. The motivation-decision model of pain (Fields 2018) states that actions are influenced by decisions between whether to approach something pleasant (a reward) or avoid something unpleasant (pain/loss). In this meta-analysis, we investigate the interaction between the reward and pain system and how their associated neurotransmitters, dopamine and opioid respectively, are connected. Given that PET radioligand studies are only able to include one radioligand at a time, it is difficult to study these interactions. By mapping more specific receptor co-activation, we hope to provide a tool to assist in the development of less potent and more targeted medications in order to help treat pain conditions while simultaneously preventing physical dependence.

One of the biggest clusters of overlap was found within striatal regions, suggesting that both dopamine and opioid receptor-mediated mechanisms are involved in modulating the perception of sensory/emotional pain. The basal ganglia circuits contain one of the highest levels of endogenous opioids and opioid receptors on the brain (McDonald & Lambert, 2005). All opioid receptor subtypes are present in the basal ganglia circuit and their net effect depends on their presynaptic or postsynaptic localization in the different structures of this circuit (Sulzer et al., 2016). Furthermore, both the midbrain VTA and nucleus accumbens receive beta-endorphin– containing projections from the arcuate nucleus and contain enkephalinergic interneurons. Both beta-endorphin and enkephalins, acting via receptors, inhibit GABA release from local inhibitory interneurons, thus facilitating dopamine release in the nucleus accumbens. D2 striatal cells modulate the indirect dopaminergic pathway, which is activated when a stimulus is less rewarding than predicted, suggesting that this is the pathway in which avoidance to an unpleasant stimulus is encoded (Bromberg-Martin, Matsumoto et al. 2010). A PET study showed that a single dose of the receptor agonist remifentanil elicited a decrease in the binding of the D2/D3 receptor radiotracer FAL in the ventral striatum, dorsal putamen, and amygdala, reflecting a release of endogenous opioids in both alcohol-dependent patients and controls (Spreckelmeyer, Paulzen et al. 2011). Our results are consistent with co-release of these neurotransmitters in these regions. Along with the existing literature (Sulzer et al. 2016, Kirkpatrick et al. 2015, Reisi et al. 2014), our results suggest that pain modulation during sensory/emotional pain conditions activates both dopamine and opioid receptors. Our results support the existing literature, in that we found the striatum to have the highest activation in both dopaminergic and opioidergic meta-analyses.

Several limitations should be noted. Our meta-analysis was intended to compare only painful or unpleasant experimental conditions. In the case of the dopamine literature, given that most work utilizing RAC used rewarding and/or pleasant experimental conditions, there is not much pain related research to include here. Therefore, while n=166 is large enough to conduct a meta-analysis, the findings might not be as robust as in our opioid meta-analysis of n=283. Given the sex differences in the neurobiology of pain, another limitation of the study involves the inability to conduct a gender analysis due to the small number of studies that provided coordinates for only male or female participants.

## 5. Conclusion

To our knowledge, this is the first powerful, data-driven meta-analysis identifying and comparing the neural correlates of opioidergic and dopaminergic transmission in PET radioligand studies of pain. The study design enabled high validity and statistical power by including 449 subjects.

The motivation for the current meta-analysis was to examine the regions in which opioid and dopamine pathways were activated during pain-related and unpleasant conditions, and consequently create a map of the distinct localization of neurotransmitters as well as their overlap. Areas that are known to be consistently activated by pain modulation are the insula, ACC, hypothalamus, PAG, rostral ventral medulla, and spinal cord (Tracey, 2010). After performing a mean analysis of opioid and dopamine binding, we found activity in these previously identified regions, but additionally were able to determine what activity was due to opioid versus dopamine neurotransmission. The significance of the resulting conjunction map (Figure 3) indicates that pain modulation is co-managed by opioid and DA receptors in a number of brain regions. Understanding the anatomical arrangement of these types of receptors may help us to further understand the interplay between the neural networks involved in pain modulation and reward. By creating maps of opioid and dopamine receptor activation, along with a map of their co-activation during pain modulation, we hope to provide data that can be used to develop targeted pharmaceuticals for patients with pain conditions. For example, animal research on drugs targeting a combination of opioid receptors have found that when targeting δ and μ-opioid receptors, the efficacy of μ-opioid receptors increases and tolerance and dependence are attenuated (Ananthan, 2006). By mapping μ-opioid receptor (CFN) and non-specific opioid receptor (DPN) activation, we hope to provide a tool to assist in the development of less potent and more targeted medications in order to help treat pain conditions while simultaneously preventing physical dependence.

## FUNDING AND DISCLOSURES

This work was supported by … ADS is supported by a VA Clinical Science Research and Development Merit Grant (I01 CX001762).

## CONFLICT OF INTEREST STATEMENT

All authors declare no conflicts of interest.

## Notes

### Competing Interest Statement

The authors have declared no competing interest.

